# Rearing temperature and fatty acid supplementation jointly affect membrane fluidity and heat tolerance in *Daphnia*

**DOI:** 10.1101/295998

**Authors:** D. Martin-Creuzburg, B.L. Coggins, D. Ebert, L.Y. Yampolsky

**Affiliations:** Limnological Institute, University of Konstanz, Mainaustrasse 252, 78464 Konstanz, Germany; Department of Biological Sciences, East Tennessee State University, Johnson City TN 37601 USA; Zoology, Department of Environmental Sciences, Zoology, University of Basel, Vesalgasse 1, 4051, Basel, Switzerland.

**Keywords:** acclimation, *Daphnia magna*, eicosapentaenoic acid, fluorescence polarization, heat tolerance, membrane fluidity, polyunsaturated fatty acids, temperature

## Abstract

The homeoviscous adaptation hypothesis states that the relative abundance of polyunsaturated fatty acids (PUFA) decreases in the membrane phospholipids of ectothermic organisms at higher temperatures to maintain vital membrane properties. We hypothesized that the well-documented reduced heat tolerance of cold-reared *Daphnia* is due to the accumulation of PUFA in their body tissues and that heat-reared *Daphnia* contain reduced amounts of PUFA even when receiving a high dietary supply of PUFA. In *Daphnia* reared at 15°C, supplementation of a PUFA-deficient food with the long-chain PUFA eicosapentaenoic acid (EPA) resulted in an increase in the relative abundance of EPA in body tissues and a decrease in heat tolerance. However, the same was observed in *Daphnia* reared at 25°C, indicating that the ability of heat-acclimated *Daphnia* to adjust EPA body concentrations is limited when exposed to high dietary EPA concentrations. *Daphnia* reared at 25°C showed the lowest change in membrane fluidity, measured as fluorescence polarization. For *Daphnia* reared at three different temperatures, thermal tolerance (time to immobility at a lethally high temperature) and increasing dietary EPA concentrations correlated with fluorescence polarization and the degree of fatty acid unsaturation. Overall, our results support the homeoviscous adaptation hypothesis by showing that cold-reared *Daphnia,* which accumulate PUFA within their tissues, are more susceptible to heat than hot-reared *Daphnia,* which contain less PUFA.

## Introduction

The ability of ectothermic organisms to cope with environmental temperature changes depends on physiological acclimation capacities and temperature-related heritable traits. Regulatory processes involved in physiological temperature acclimation include the up-regulation of heat shock proteins that reduce protein denaturation (Richter et al. 2010), up-regulation of antioxidant pathways that limit oxidative damage at high temperatures (Lushchak 2011), and membrane remodeling that allows membranes to maintain optimal viscosity over a range of temperatures (Sinensky 1974; Hazel and Williams 1990; Hazel 1995). The homeoviscous adaptation hypothesis states that ectothermic organisms adjust the proportion of mono-or polyunsaturated fatty acids (PUFA) in their membrane phospholipids to the prevailing temperatures (Hazel and Prosser 1974; Hazel 1995; Ballweg and Ernst 2017), the highest degree of fatty acid unsaturation expected at the lowest temperature. Long-chain PUFA, i.e. those with 20 or more carbon atoms, often accumulate in cold-exposed and winter-active ectotherms, suggesting that they play a role in the acclimation to low temperatures (Farkas 1979; Schlechtriem et al. 2006; Smyntek et al. 2008).

With few exceptions, animals cannot synthesize long-chain PUFA *de novo* and have only limited abilities to synthesize long-chain PUFA from dietary C18 PUFA precursors. Thus, most animals rely on a dietary supply of these essential compounds (Cunnane 2000; Leonard et al. 2004; Schlechtriem et al. 2006). The PUFA requirements of ectotherms have been intensively studied using the freshwater crustacean *Daphnia* as a model. A deficiency in dietary PUFA, especially in the C20 PUFA eicosapentaenoic acid (EPA), has been shown to severely constrain *Daphnia’s* growth and reproduction (von Elert 2002; Becker and Boersma 2003; Martin-Creuzburg et al. 2009). Experimental evidence suggests that PUFA requirements increase with decreasing temperature (Martin-Creuzburg et al. 2012; Sperfeld and Wacker 2011; 2012). A high dietary PUFA supply at low temperatures may increase the incorporation of PUFA into membrane phospholipids, thus increasing membrane fluidity and allowing *Daphnia* to cope with lower water temperatures. At high temperatures, however, high dietary PUFA supply may also result in the increased incorporation of PUFA into membrane phospholipids, which may be detrimental for maintaining important membrane properties and thus may reduce *Daphnia’s* tolerance to heat. Besides that, PUFA are primary targets of lipid peroxidation (LPO), resulting in the formation of toxic degradation products (Gueraud et al. 2010; Lushchak 2011). The effects of temperature on lipid peroxidation are controversial, but most studies have found increased oxidative stress at higher environmental temperatures (Heise et al. 2006; Bagnyukova et al. 2007; Lushchak 2011), supporting the idea that a high dietary PUFA supply is detrimental at high temperatures.

Despite the well-documented increase in heat tolerance in *Daphnia* raised at high temperatures (Paul et al. 2004; Zeis et al. 2004; Williams et al. 2012; Yampolsky et al. 2014), as well as a parallel decrease in the relative abundance of PUFA in body tissues (Masclaux et al. 2012), a causative relationship between PUFA abundance and heat tolerance has not been proven. Furthermore, it is unclear if membrane fluidity is a crucial aspect of this relationship. Here, we analyzed the joint effects of environmental temperature and dietary EPA availability on fatty acid composition, membrane fluidity, and heat tolerance in three clones of *Daphnia magna.* Membrane fluidity was measured as fluorescence polarization (FP), which quantifies the amount of order in membranes and other hydrophobic cellular components and thus is a measure of membrane integrity (Shinitzky and Barenholz 1978; Kushnareva 2009; Dawaliby et al. 2016; Ballweg and Ernst 2017). FP decreases (and thus membrane fluidity increases) with increasing assay temperature as the membranes melt. It has been demonstrated that membrane fluidity in vivo is held constant across temperatures, presumably by homeoviscous adaptation mechanisms (Hazel and Prosser 1974; Hazel and Williams 1990; Hazel 1995; Hochachka and Somero 2002; Ballweg and Ernst 2017). Here, we tested the hypothesis that dietary EPA supplementation is linked to higher membrane fluidity, thereby reducing heat tolerance in the freshwater crustacean *Daphnia magna.*

## Materials and Methods

### Liposome preparation

Liposome stock suspensions were prepared using 3 mg 1-palmitoyl-2-oleoyl-phosphatidylglycerol (POPG) and 7 mg 1-palmitoyl-2-oleoyl-phosphatidylcholin (POPC; Lipoid, Germany) dissolved in aliquots of ethanol as described in Martin-Creuzburg et al. (2009). EPA-containing liposomes were prepared by adding 3.33 mg of EPA (eicosapentaenoic acid, 20:5n-3; Sigma-Aldrich) from lipid stock solutions in ethanol (2.5 mg mL^−1^). The resulting solutions were dried using a rotary evaporator, dissolved in 10 mL buffer (20 mmol I^−1^ NaP,, 150 mmol |^−1^ NaCI, pH 7.0), and vigorously sonicated in an ultrasonic bath (2 min). Excess free EPA was removed by washing the liposomes in fresh buffer using an ultra-speed centrifuge (150,000 g, 90 min, 4°C). The liposome stock suspensions were sonicated again (2 min) before liposomes were added to the experimental jars. The liposome stock suspensions contained approximately 1×10^6^ liposomes mL^−1^ with a mean diameter of 3.4 μm.

### Dietary manipulation

We used three *Daphnia magna* clones originating from geographically distinct locations in the UK, Hungary, and Israel (Supplementary Table 1). Juveniles less than 24 h old were collected from 400-mL jars containing eight females, which were fed with 250,000 cells of *Scenedesmus obliquus* per mL every other day (four replicate jars per clone). Juveniles were placed into 100-mL jars (two individuals per jar) containing 80 mL of ADaM, artificial *Daphnia* medium (Klüttgen et al. 1994), and reared in either 15°C, 20°C or 25°C incubators. We collected neonates from the same mothers for 10 days, allowing 5-7 days between each temperature increment, so that the *Daphnia* at each of the three temperatures reached maturity at approximately the same time. The animals were fed S. *obliquus* (to the final concentration of 50,000 cells/mL) supplemented with 50 μL of one of four liposome suspensions: either 100% control liposomes; 2/3 control + 1/3 EPA-containing liposomes; 1/3 control + 2/3 EPA-containing liposomes, or 100% EPA-containing liposomes. The 100% EPA-supplemented liposome diet corresponded to 1.51 ± 0.09 μg EPA per *Daphnia* per day. *S*. *obliquus* was used because it does not contain PUFA with more than 18 carbon atoms. Food was added every other day and water was changed once every four days. The position and orientation of trays within the incubator were rotated after each feeding. In the cases of mortality of one of the two individuals in a jar, individuals from two identical jars were combined to maintain two individuals per jar. One individual from each jar was used for FP measurement and the other one for either thermal tolerance testing or fatty acid analysis. Two individuals from the same clone and treatment, but from different jars, were combined to form a single replicate for the fatty acid analysis. Four replicates were created for each combination of temperature, diet and clone, resulting in four replicates for determination of fluorescence polarization, two for heat tolerance measurement, and two for fatty acid analysis. However, due to mortality, the average numbers of replicates per clone, per treatment were 3.6 for FP and 1.8 for heat tolerance testing. In addition, three of the fatty acid analysis replicates consisted of one rather than two individuals. Females carrying their 2^nd^ or 3^rd^ clutch were used in each of these measurements.

The individuals used for fatty acid analysis were starved for 24 hours, incubated in a dextran beads emulsion (Sephadex, 30 to 50 μm diameter) for another 24 hours to clear the guts from algae and liposomes, and stored at −80°C in pre-weighed aluminum boats. Prior to fatty acid analysis, all samples were freeze-dried and weighed for dry mass determination.

### Fatty acid analysis

For the analysis of fatty acids in the liposomes and animal tissues, aliquots of the liposome suspensions or two animals, respectively, were deposited in 7 mL of a mixture of dichloromethane and methanol (2: 1, v: v), sonicated vigorously, and stored at −20°C overnight. Total lipids were extracted three times with dichloromethane: methanol (2: 1, v: v). Pooled cell-free extracts were evaporated to dryness under a stream of nitrogen. The lipid extracts were transesterified with 3 mol L^−1^ methanolic HCI (60°C, 15 min), and the fatty acid methyl esters (FAME) were extracted three times with 2 mL of iso-hexane. The FAME-containing fraction was evaporated to dryness under nitrogen and resuspended in a volume of 10-20 μL iso-hexane. FAME were analyzed by gas chromatography on a HP 6890 gas chromatograph (GC) equipped with a flame ionization detector (FID) and a DB-225 (J&W Scientific, 30 m × 0.25 mm inner diameter × 0.25 μm film) capillary column. Details of GC configurations are given elsewhere (Martin-Creuzburg et al. 2010). Fatty acids were quantified by comparison to an internal standard (C23:0 ME) of known concentration, considering response curves determined previously with fatty acid standards (Sigma-Aldrich). Fatty acids were identified by their retention times and their mass spectra, which were recorded with a gas chromatograph-mass spectrometer (GC-MS; Agilent Technologies, 5975C inert MSD) equipped with a fused-silica capillary column (DB-225MS, J&W Scientific; GC configurations as described for FID). Mass spectra were recorded between m/z 50 and 600 in the electron ionization (EI) mode. The limit of quantification was 10 ng of fatty acid. To calculate relative abundances, the absolute amount of each fatty acid was related to either the animal’s dry mass or to total FA content. Relative abundance of each of the FAs, except EPA itself, was calculated excluding EPA from the total. To characterize the degree of fatty acid unsaturation in samples, we calculated the unsaturation index as 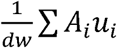 where *dw* is the dry weight of the sample (mg), *A*_*i,*_ the abundance (ng), and *u*_*i*_, the unsaturation level (number of double bonds) of *i*-th fatty acid.

### Fluorescence polarization

Fluorescence polarization (FP) is believed to quantify the amount of order in cell membranes and other hydrophobic cellular components (Shinitzky and Barenholz 1978; Dawaliby et al. 2016; Ballweg and Ernst 2017). FP is reciprocally related to membrane fluidity: as membrane fluidization increases the mobility of the fluorophore dye, it decreases the intensity of the emitted parallel component. FP is thus calculated as the ratio between fluorescence intensities measured with parallel and perpendicular position of the emission filter.

To quantify FP, individual *Daphnia* were homogenized in 100 μL of HEPES buffer (pH 7.4), containing 10 mM KCl, 20 mM HEPES, 1 mM EGTA, 2 mM EDTA, 250 mM sucrose, 1 mM DTT, 5mL/L protein inhibitor cocktail III, and 10 μΜ DPH fluorophore, on a MP plate homogenizer, using 0.15 mm glass beads, centrifuged for 4 min at 4000 rpm at 4°C, and incubated for 30 minutes at room temperature. Subsequently, 40 μΙ aliquots of the samples were transferred into a black low-absorption, low volume 384-well plate, which was equilibrated to temperatures from 10 to 45°C with a 5°C increment in a thermocycler. The plates were incubated at each target temperature for 5 minutes (estimated by measurement of temperature in several blank wells). FP was then measured on a Biotek Synergy H1 plate-reader (equilibrated to each temperature) with 360 nm excitation and 460 nm emission filters. Gain and optics positions were calibrated according to manufacturer’s instructions. The temperature of several blank wells was measured before and after the fluorescence measurements, and the average was recorded. This resulted in slight deviations from the desired temperatures — upward from the target temperature for temperatures below room temperature, and downward for temperatures above room temperature. FP was calculated as

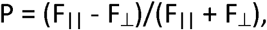

where F_||_ and F_⊥_ are parallel and perpendicular fluorescence measurements, respectively. To assess the assay temperature-independent magnitude of FP for each well, a linear regression was calculated over the assay temperate range from 20 to 45°C and evaluated at 30°C. This temperature is close to the midpoint of the assay temperature range, thus providing the best statistical estimate of the vertical position of the regression line. This temperature is also close to the upper long-term survival limit for typical *D. magna* clones, thus possibly reflecting a biologically important threshold. This quantity is referred to as FP_30_. The remainder of the sample was used for protein content (Bradford 1974) and lipid peroxidation measurements (as described in Coggins et al. 2017).

### Heat tolerance

Time until immobilization (T_imm_) at 37°C (Yampolsky et al. 2014) was used as the measure of acute temperature tolerance. Briefly, individual females were placed in cylindrical glass vials containing 50 mL of ADaM at 20°C and transferred into a 37°C water bath resulting in a 10-15-minute temperature ramp. We monitored swimming ability every 1-3 minutes until the *Daphnia* was no longer capable to sustain itself in the water column. This time was recorded as T_imm_, and its logarithm was used in the analysis for the sake of normality.

### Statistical analysis

Data were analyzed by applying general linear mixed models using JMP 10 (JMP, 2012) with rearing temperature and EPA supplement as continuous fixed effects and clones as a random effect. The alternative analysis with temperature and EPA supplement as categorical effects yielded identical results and is not reported here (with one exception, see below). In the joint analysis of FP_30_, FA composition and T_imm_, we used clonal means for each combination of rearing temperature and EPA supplement. To untangle the effects of FP_30_ and unsaturation index on T_imm_ from the general effects of rearing temperature and EPA supplement, we first calculated the residuals of the log-transformed T_imm_ values over the rearing temperature and EPA supplements (and their interaction) as continuous effects influencing T_imm_. We then used linear regressions of these residuals on either FP_30_ or unsaturation index. Principal component analysis on covariances of fatty acid relative abundance was done in JMP 10 (JMP 2012).

## Results

### Fluorescence polarization

Fluorescence polarization decreased linearly with the assay temperature between 20 and 45°C (supplementary Fig. S1, Table 1). FP_30_ was highest at the highest rearing temperature, consistent with the homeoviscous adaptation model (Fig. 1; Table 1). FP_30_ decreased with rearing temperature for low dietary EPA concentrations (0 and 33%), but not for high dietary EPA concentrations (66 and 100%; Fig. 1, supplementary Fig. S1). The rearing temperature } EPA interaction was not significant in the regression analysis (Table 1) but was significant if rearing temperature and EPA interaction were treated as categorical effects (df_num = 3, df_den = 113, F = 4.59, P < 0.001). There were no differences among acclimation temperatures or EPA treatments in Bradford protein content (data not presented), indicating that the observed differences in FP are not an artifact of sample concentration or protein denaturation.

**Fig. 1:**
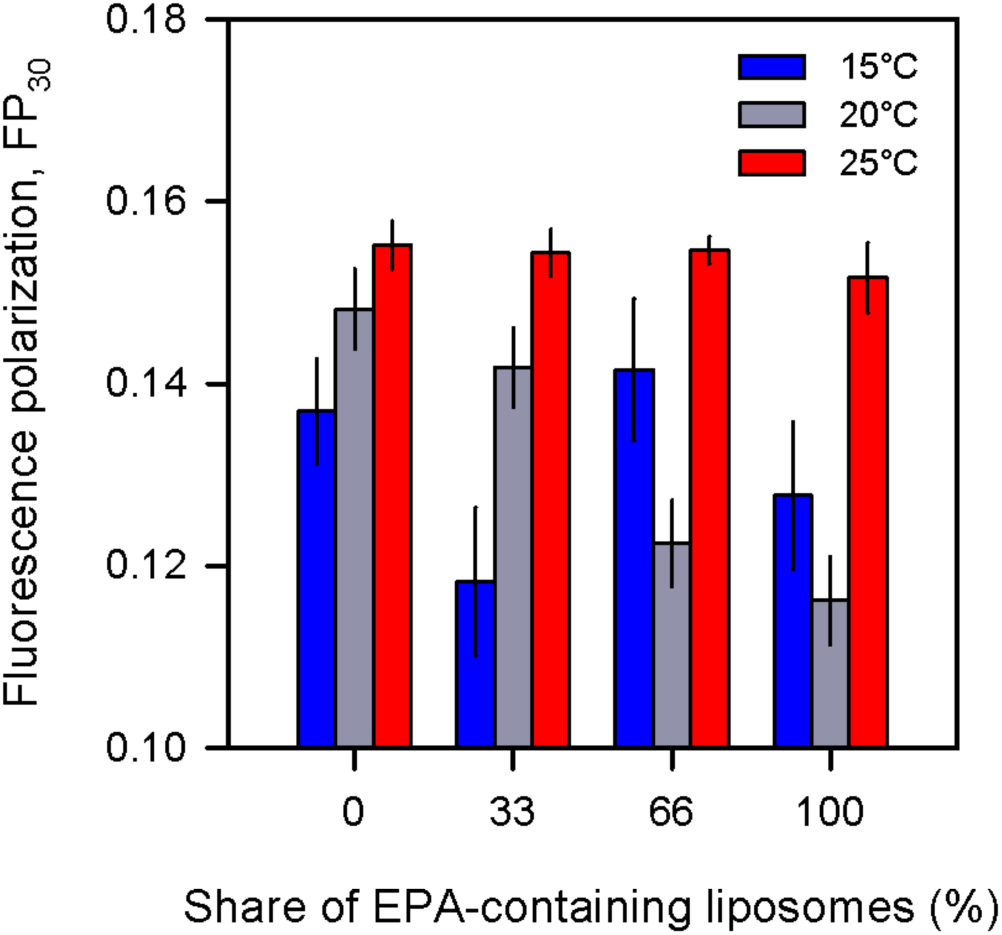
FP_30_ values for *Daphnia magna* reared at 15^°^C (blue), 20°C (gray), and 25 °C (red) and fed dietary supplements with different EPA concentrations, i.e. *S*. *obliquus* supplemented with 0, 33, 66 or 100% EPA-enriched liposomes. Data are means of 6-12 replicates per temperature/EPA combination; vertical bars represent standard errors.

**Table 1:** Fixed effects test of FP_30_ as a function of rearing temperature (T) and EPA

| Source | DF | SS | MS | F Ratio | Prob > F |
| --- | --- | --- | --- | --- | --- |
| T | 1 | 0.0097 | 0.0097 | 32.47 | 8.4E-8 |
| EPA | 1 | 0.0040 | 0.0040 | 13.29 | 0.0004 |
| T*EPA | 1 | 0.00003 | 0.00003 | 0.12 | 0.73 |
| Clone | 2 | 0.0035 | 0.0017 | 5.78 | 0.004 |
| Error | 123 | 0.0366 | 0.0003 |  |  |

### Fatty acid analysis

The two dominant C16 fatty acids, C16:0 and C16:1n-7, did not show consistent patterns in response to temperature or EPA treatment (Fig. 2A and B, Table 2 and Table S2). The dominant monounsaturated fatty acid C18:1n-9 increased with dietary EPA supply (Fig. 2C), whereas C18 PUFA, linoleic acid (C18:2n-6) and α-linolenic acid (C18:3n-3) decreased at all temperatures with increasing EPA supplementation (Fig. 2D,E). Moreover, whereas α-linolenic acid (C18:3n-3) decreased in relative abundance with increasing rearing temperature, linoleic acid (C18:2n-6) increased in *Daphnia* reared at 25°C(Fig. 2D,E; Table 2). Taken together, both temperature and EPA treatment profoundly influenced the balance between the two major PUFA in body tissues. None of the temperature × diet interactions approached significance, suggesting that the effects of EPA availability and temperature are independent from each other.

**Fig. 2:**
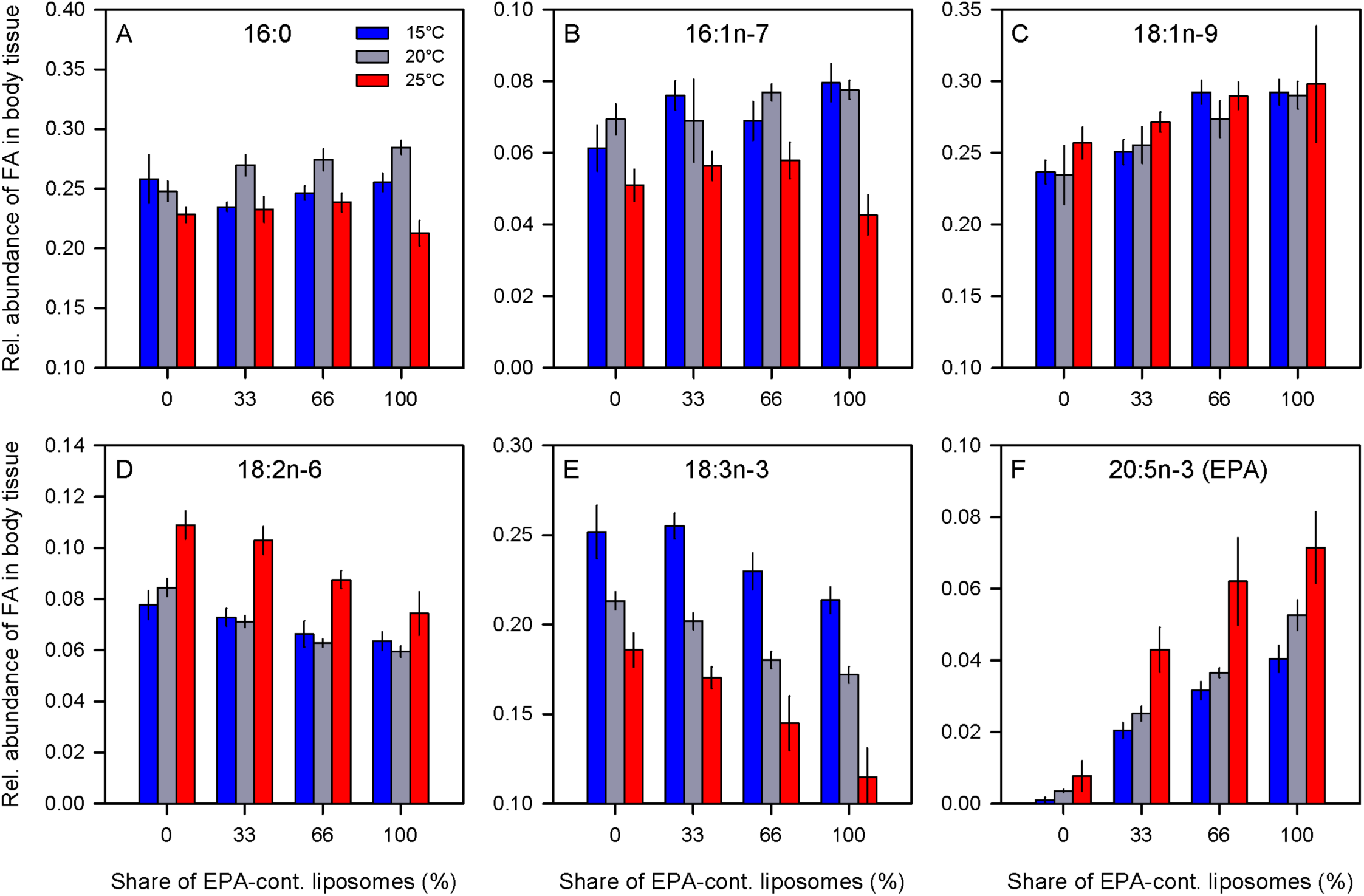
Relative abundance (normalized to total FA content) of fatty acids in *Daphnia magna* body tissue. The figure depicts fatty acids that were significantly influenced by rearing temperature or dietary EPA supply and represented at least 5% of total fatty acids (A-E; shown in bold in Table 1) and EPA (20:5n-3; F). Data represent means of n = 6 per EPA/temperature combination and standard errors.

**Table 2:** Fatty acid responses to rearing temperature and dietary EPA supply. The table lists only fatty acids for which the rearing temperature (T) and the dietary EPA supply (EPA) were still significant after sequential Bonferroni correction (P < 0.01) and which had an average share of total fatty acids of at least 0.01 (bold: at least 0.05; see Fig. 2). None of the interaction terms (TxEPA) were significant. The T and EPA column depicts increasing (Up) or decreasing (Down) shares of a particular fatty acids relative to total fatty acids, excluding the supplemented EPA (ANOVA results are presented in Table S2).

| Fatty acid | T | EPA |
| --- | --- | --- |
| C14:0 | Down | - |
| **C16:0** | **Down** | - |
| C18:0 | Up | - |
| **C16:1n-7** | **Down** | - |
| C18:1n-7 | Up |  |
| **C18:1n-9/n-12** | - | **Up** |
| **C18:2n-6 (Linoleic acid)** | **Up** | **Down** |
| **C18:3n-3 (α-Linolenic acid)** | **Down** | **Down** |
| C18:4n-3 | Down | Down |
| C20:5n-3 (EPA) | Up | Up |

Supplementation with EPA resulted in increasing concentrations of EPA in tissue extracts and, regardless of the dietary availability of EPA, the relative abundance of this PUFA increased with rearing temperature (Fig. 2F). As total fatty acid content was the lowest at the highest acclimation temperature (supplementary Fig. S2B), the temperature effect depended on the normalization type, i.e. whether the fatty acid data were related to the total fatty acid content or to the dry mass of the samples. However, we observed an increase in EPA body concentrations with dietary EPA supply with both types of normalization (Fig. S2A). On the other hand, normalization by dry weight eliminated the unexpected high EPA concentrations in animals reared at 25°C (Fig. 3F), due to the lowest total fatty acid accumulation at this temperature (Fig. S1B). The highest dry mass-normalized EPA concentration was observed at the intermediate (20°C) acclimation temperature (Fig. S2A).

**Fig. 3:**
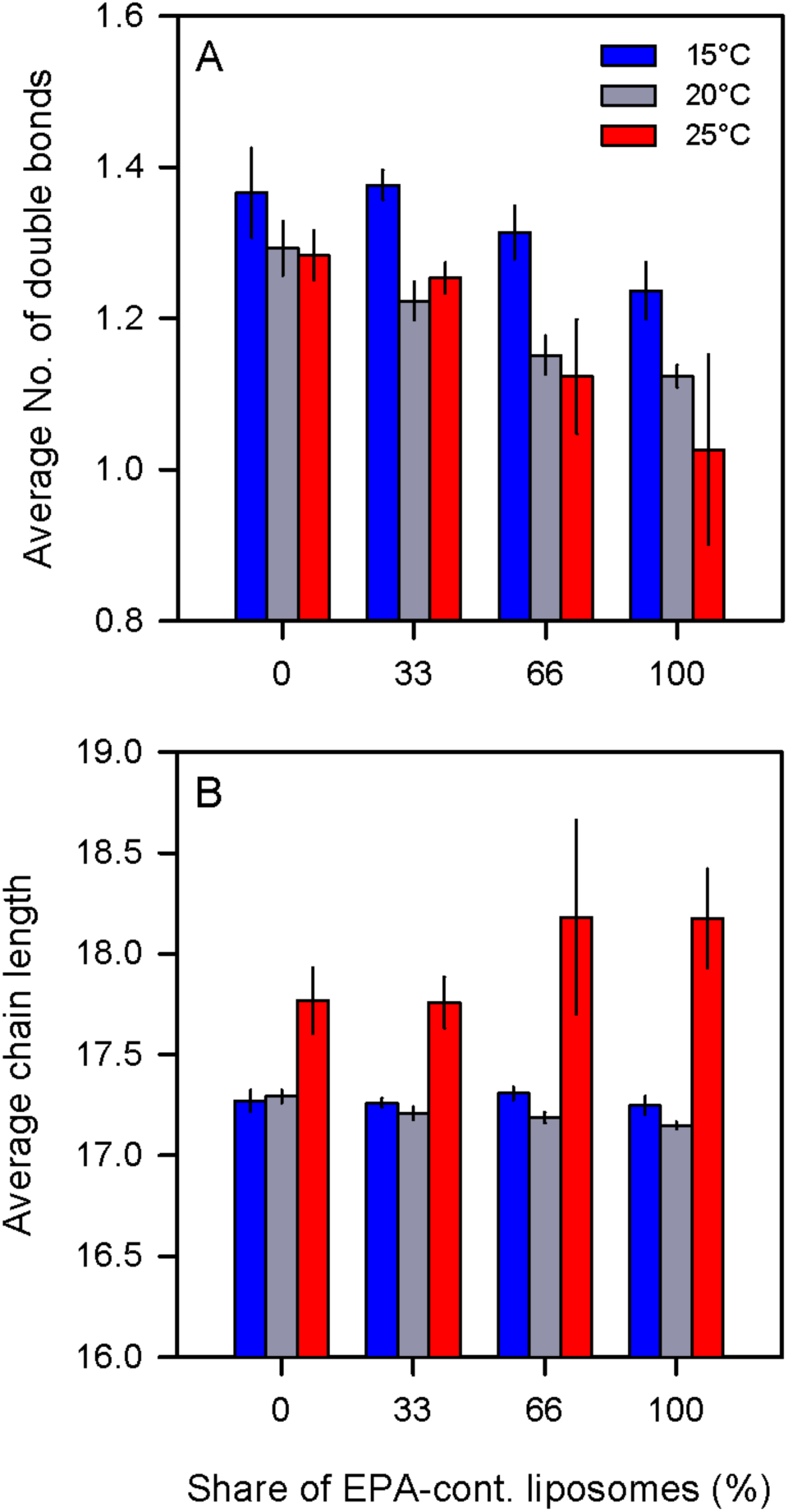
Average number of double bonds (A) and average chain length (B) of fatty acids, excluding EPA, detected in *Daphnia magna* reared at different temperatures and dietary EPA concentrations. Data represent means of n = 6 per EPA/temperature combination and standard errors.

Changes in fatty acid composition (excluding EPA itself) were associated with changes in the number of double bonds and carbon atoms within fatty acid chains (Fig. 3). Rearing temperature had a significant effect on chain length (df_num_ = 2, df_den_ = 56, F = 5.11; P < 0.01; Fig. 3B) but not on the number of double bonds (df_num_ = 2, df_den_ = 56, F = 1.00, P > 0.37). Inversely, dietary EPA supply had an effect on the number of double bonds (df_num_ = 3, df_den_ = 56, F = 9.55, P < 0.001) but not on the chain length (df_num_= 3, df_den_ = 56, F = 0.95, P > 0.42). The temperature x EPA interaction was not significant in either analysis.

Principal component analysis of the relative abundances of all fatty acids (EPA excluded from the denominator) revealed a strong separation between the effects of temperature and EPA treatments along the first two principal components (PC), explaining 41.6% and 30.1% of variance (Fig. 4). The relative abundance of saturated fatty acids and PUFA had a significant leverage on the PC1, which significantly separated the three acclimation temperatures, as well as the four diet regimes (Fig. 4).

**Fig. 4:**
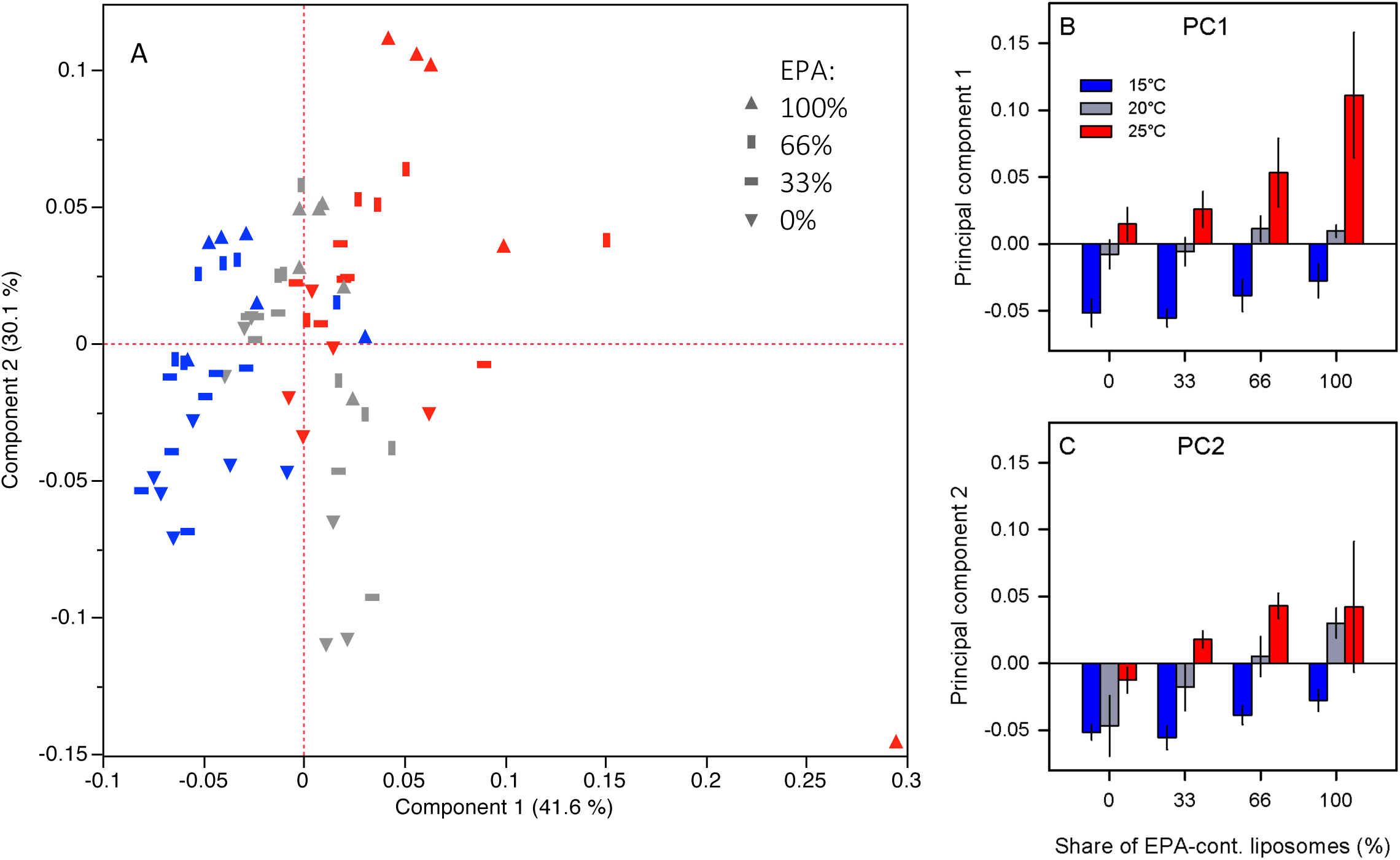
Principal component analysis (A) of relative fatty acid abundances (EPA excluded from the denominator). Symbols represent levels of dietary EPA, i.e. 0, 33, 66, and 100 % of EPA-containing liposomes; colors represent rearing temperatures, i.e. 15°C (blue), 20°C (gray) and 25°C (red). Principal components 1 and 2 are also presented as a function of rearing temperature and EPA content (B-C).

### Temperature tolerance

Rearing temperature and dietary EPA supply had significant effects on heat tolerance measured as time until immobilization (T_imm_) at 37°C (Fig. 5, Table 3), whereas the T × EPA interaction was not significant. At the highest dietary EPA supply, T_imm_ was significantly reduced in *Daphnia* reared at 15°C and 25°C (Fig. 5).

**Table 3:** Fixed effects test of rearing temperature (T) and dietary EPA supply (EPA) on log-transformed T_imm_ with clones as a random effect.

| Source | DF | SS | MS | F Ratio | Prob > F |
| --- | --- | --- | --- | --- | --- |
| T | 1 | 14.659 | 14.659 | 103.49 | 1.6E-14 |
| EPA | 1 | 1.510 | 1.510 | 10.66 | 0.0018 |
| T*EPA | 1 | 0.0015 | 0.0015 | 0.01 | 0.92 |
| Clone | 2 | 1.761 | 0.881 | 6.21 | 0.004 |
| Error | 58 | 8.215 | 0.142 |  |  |

**Fig. 5:**
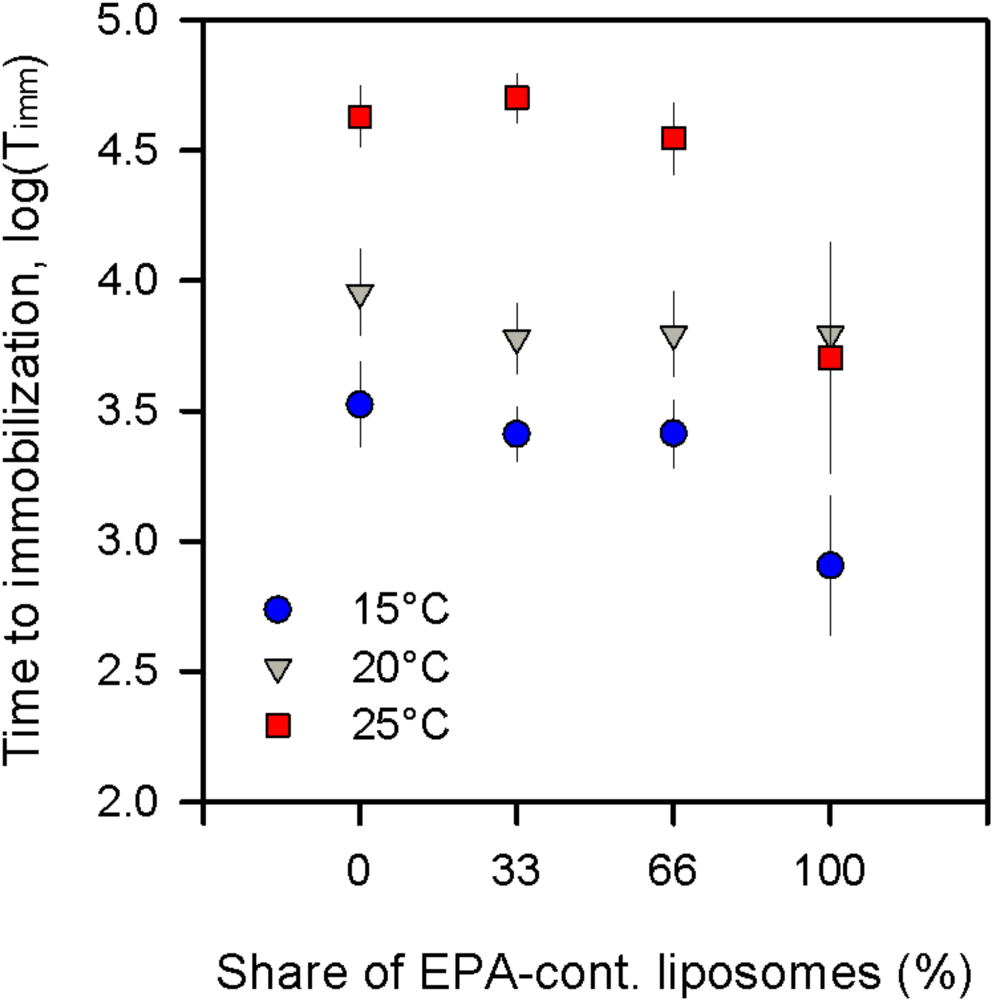
Effects of rearing temperature and dietary EPA supply on acute high temperature tolerance represented as time to immobilization (log-transformed **T_imm_).**

### Analysis of clonal means

Because FP_30_, T_imm_, and FA composition were not measured in the same individuals, their joint analysis was only possible by correlating mean values from individuals matched by clone, rearing temperature and EPA supplementation. As expected, clonal means of the unsaturation index showed a negative effect on clonal means of FP_30_ (Fig. 6A). The unsaturation index and FP_30_ were correlated with T_imm_ (negatively and positively, respectively) across rearing temperatures and dietary EPA concentrations (Fig. 6B,C). However, it was not possible to untangle the effects of unsaturation index or FP_30_on T_imm_ from the overall effect of rearing temperature. The residuals from a multiple regression of T_imm_ on rearing temperature showed no correlation with the unsaturation index (regression coefficient ± SE was 0.0003 ± 0.0117, nor with FP_30_ (regression coefficient ± SE = −0.995 3.961).

**Fig. 6:**
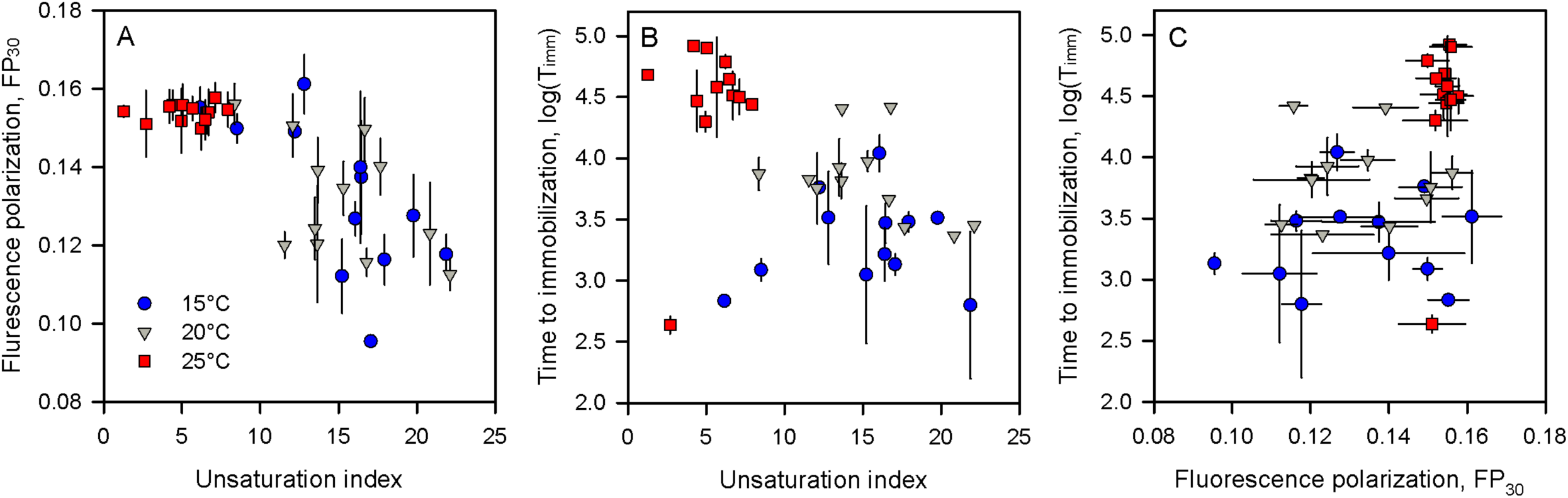
Relationships between clonal means of the level of unsaturation (unsaturation index, calculated as weighted average of fatty acid abundances weighted by the number of double bonds in each fatty acid; A and B), fluorescence polarization (FP_30_; A and C), and time until immobilization (T_imm_; B and C) measured in individuals matched by rearing temperature, dietary EPA supply, and clone.

## Discussion

In line with the homeoviscous adaptation hypothesis (Hazel and Williams 1990; Hazel 1995; Hochachka and Somero 2002; Ballweg and Ernst 2017), we show that membrane fluidity decreases (i.e. FP increases) with increasing rearing temperature (Fig. 1; Table 1) and increases (i.e. FP decreases) with the degree of fatty acid unsaturation (Fig. 6A). This result suggests that *Daphnia* are capable of adjusting their membrane lipid composition to higher temperatures by reducing the degree of phospholipid fatty acid unsaturation. Consistent with previous findings (Zeis et al. 2004; Yampolsky et al. 2014), T_imm_ was positively correlated with rearing temperature (Fig. 5). When the clonal means of FP_30_ and T_imm_ were analyzed together across treatment combinations, the observed effect of rearing temperature was mirrored by the correlation between FP_30_, unsaturation index, and T_imm_ (Fig. 6). Because we were unable to untangle the overall effects of rearing temperature and dietary EPA supply from the specific effects of unsaturation index or FP_30_ on T_imm_, we cannot claim that the effect of rearing temperature on heat tolerance can be explained statistically by lipid composition and membrane fluidity. However, our results are consistent with the hypothesis that the well-known effect of rearing temperature on heat tolerance occurs through the adjustment of the level of FA unsaturation, which in turn affects membrane fluidity.

The relationship between rearing temperature, PUFA availability and PUFA accumulation in tissues is not simple. Contrary to the prediction that organisms reared at high temperatures should contain lower amounts of PUFA, the relative abundance of EPA in *Daphnia* reared at 25°C increased significantly with increasing dietary EPA availability, both relative to the total fatty acid content and relative to the dry weight (Fig. 2, Fig. S2A). The average number of double bonds in the hydrocarbon chains of fatty acids and the relative abundance of α-linolenic acid decreased with acclimation temperature, which is consistent with the homeoviscous adaptation hypothesis (Fig. 3A, Table 2). However, the abundance of other FA did not follow this trend. It appears that *Daphnia* cannot avoid EPA accumulation at high dietary EPA supply. This EPA accumulation may be the proximate cause of the reduced temperature tolerance of *Daphnia* reared on the EPA supplemented diets at the highest temperature (Fig. 5).

The dependence of temperature tolerance on rearing conditions observed in this study did not always follow predicted linear relationships; for example, T_imm_ was significantly lower at high dietary EPA supply in 15°C- and 25°C-reared *Daphnia* but did not change with increasing dietary EPA supply in 20°C-reared *Daphnia.* Thus, this dependence is likely more complex than the analysis of FA composition allows us to detect here. For example, membrane fluidity can also be modulated by changing the relative proportions of phospholipid classes and of other membrane components, such as sterols (Dufourc 2008; Fajardo et al. 2011; Dawaliby et al. 2016). Differences in relative abundances of these membrane components, not measured in this study, may explain why the FP_30_ values decreased with dietary EPA supply only in *Daphnia* reared at the intermediate temperature of 20°C (Fig. 1) and why the expected decrease in heat tolerance with EPA supplementation was observed only in *Daphnia* reared at 15°C- and 25°C. The assay we used to measure FP_30_ was based on whole-body extracts and thus we did not specifically measure fatty acids incorporated into membrane phospholipids or, for example, storage triglycerides. The observed EPA accumulation at higher temperature may therefore not contradict the homeoviscous adaptation hypothesis in the sense that EPA was not accumulating in membranes and thus not affecting their fluidity, but was rather stored in lipid droplets or reproduction-related tissues.

In *Daphnia,* EPA is critical for growth and reproduction (von Elert 2002; Becker and Boersma 2003; Martin-Creuzburg et al. 2009; Schlotz et al. 2012), and its allocation into reproductive tissues may increase with acclimation temperature as overall resources become less abundant due to increased metabolic costs. The decrease in relative abundance of α-linolenic acid in body tissues with acclimation temperature and dietary EPA supply (Fig. 2) may result from a reduced dietary absorption of α-linolenic acid when EPA is abundant. However, dietary EPA availability cannot fully explain the inverse trends in acclimation temperature response for the relative abundances of linoleic and α-linolenic acid because these trends were also observed in *Daphnia* that received no dietary EPA supplement (Fig. 2). This observation is consistent with earlier findings (Coggins et al. 2017), indicating that most of the decrease in PUFA abundance at high acclimation temperature is due to the decrease of phospholipids with one or both fatty acid chains represented by 18:3 fatty acids (Coggins et al. 2017). It is possible that *Daphnia* do not modulate the relative abundance of 18:2 PUFA as predicted by the homeoviscous adaptation hypothesis. Although not large in magnitude, the increase in relative abundance of linoleic acid with temperature is statistically significant (Fig. 2, Table 2) and may reflect a by-product of adaptive changes in expression of PUFA pathway enzymes, for example, differential expression of paralogs and/or isoforms of Δ3- or Δ6-desaturases.

## Conclusions

In line with the homeoviscous adaptation hypothesis, this study found that dietary EPA supplementation decreased heat tolerance in *Daphnia,* as EPA simultaneously increased in body tissues. As these effects are observed in *Daphnia* raised at both low and high temperatures, the prediction of low PUFA accumulation at high temperatures is not supported by our study. *Daphnia* may have limited abilities to adjust the PUFA composition of body tissues when EPA is abundant, which may be detrimental at higher temperatures. Nevertheless, temperature-related changes in body fatty acid composition were observed, including decreased levels of unsaturation, increased average chain lengths, and changes in relative abundance of n-6 (linoleic acid) and n-3 (α-linolenic acid) PUFA with increasing temperature. Consistent with the homeoviscous adaptation hypothesis, the unsaturation index and membrane fluidity (FP_30_) correlated with heat tolerance (T_imm_) across rearing conditions, suggesting that heat tolerance is mediated via environmental effects (food and ambient temperature) on membrane fluidity.

## Acknowledgements

We are grateful to Jürgen Hottinger, Leonie Seefeldt, and Petra Merkel for laboratory assistance and Eric von Elert and Aruna Kilaru for valuable discussions. This study was supported by the Swiss National Foundation.

## Data availability

All data are available in Supplementary Materials and on Dryad (datadryad.org).

